# Effects of Ca^2+^ on encystment and growth in *Scrippsiella trochoidea*

**DOI:** 10.1101/2020.12.14.422664

**Authors:** Zhifu Wang, Weihua Feng, Jing Cao, Haifeng Zhang, Dongrong Zhang, Jian Qian, Heng Tao Xu, Zhe Hao

## Abstract

Cysts serve as a seed source for the initiation and recurrence of a harmful algal bloom (HAB) caused by dinoflagellates. And the influence of calcium on cyst formation has been relatively understudied. In the present study, we investigated the effects of calcium (Ca^2+^) on the growth and encystment of *Scrippsiella trochoidea.* We incubated *S. trochoidea* in modified f/2 media in flasks which were divided into five groups and treated with different Ca^2+^ concentrations (0, 0.2, 0.4, 0.6, and 0.8 g·L^−1^). We revealed that cell density increased with increasing Ca^2+^ concentrations; however, cell density was reduced when Ca^2+^ concentrations exceeded 0.2 g·mL^−1^. Additionally, the number of cysts and the cyst formation rate similarly increased as Ca^2+^ concentrations increased, but these were reduced when Ca^2+^ concentrations exceeded 0.4 g·mL^−1^. Lastly, *S. trochoidea* absorbed Ca^2+^ from the water when cysts were formed and under high Ca^2+^ concentrations, more calcareous thorn cysts formed.

## Introduction

During recent decades, the coastal waters of China seas have experienced many harmful algal blooms (HABs) caused by dinoflagellates(Tang et al. 2016). Many dinoflagellates generate cysts during their life cycle(Blackburn S. I et al. 2005), which play an important role in promoting HABs (Figueroa R. I et al. 2010). These cysts are usually associated with genetic recombination, maintenance, termination, and recurrence of blooms. Further, they facilitate dinoflagellate survival under unfavorable environmental conditions by protecting against viruses, grazers, and parasites, as well as by promoting population expansion (Tang et al. 2012). *Scrippsiella trochoidea* is a cosmopolitan bloom-forming dinoflagellate species that can grow well in a narrow range of temperatures, with a notable tolerance of temperatures as low as 10°C (Wang et al. 2007). *S. trochoidea* easily forms cysts when surrounding environmental conditions become unsuitable for its survival; thus, *S. trochoidea* serves as a model organism for examining dinoflagellate cysts and their role in promoting HABs. *S. trochoidea* has been reported in the USA and Japan (Ishikawa A et al. 1996) and its resting cysts represent the dominant species in Chinese coastal sediments, especially in Daya and Shenzhen Bays in the South China Sea (Wang et al. 2004).

Encystment is related to factors such as aging of cultures, nutrient stress, unfavorable light intensity, temperature changes, and bacterial attacks(Tang et al. 2012). Most research on dinoflagellate encystment has aimed at describing their life history. Despite many studies which have focused on factors influencing dinoflagellate encystment, little is known about the possible effects of metal ions on *S. trochoidea* encystment and growth. *S. trochoidea* cysts are egg or oval shaped, and often covered with calcareous thorns; further, they typically contain one or two red bodies (Cho et al. 2001). In our previous study, we identified two main cyst shapes, calcareous thorn cysts and smooth surface cysts, and we further revealed that these two cyst shapes occur in different ratios under different conditions(Wang et al. 2014).

Previous studies have indicated that the most common culture manipulation which induces sexuality in autotrophic species is nutrient starvation(István Grigorszky et al.2006). However, despite being a necessary trace element of dinoflagellates, the influences of calcium (Ca^2+^) on *S. trochoidea* growth and encystment remain understudied. Thus, will further understanding the role of calcium in cyst formation in this species be important to further understanding and mitigating HABs. Therefore, in the current study, we explored the effects of Ca^2+^ on *S. trochoidea* growth to determine its role in cyst formation; we further attempted to analyze the relationship between Ca^2+^ concentration and the proportion of different *S. trochoidea* cysts.

## Materials and Methods

The clonal and axenic strains of *S. trochoidea* used in this experiment were cultured at the Second Institute of Oceanography, Ministry of Natural Resources in China. Aged seawater was filtered through a 0.45 μm pore size cellulose nitrate membrane filter and autoclaved at 125°C for 30 min. Cultures were grown at temperatures of 25 ± 1°C, similar to sea surface temperatures in the East China Sea. The cultures were grown under 300-500 lx of cool white fluorescent illumination on a 12 h light/12 h dark cycle with a salinity of 30%. To prepare bulk cultures, these growth procedures were scaled up in 3 L flasks. To prevent clumping, the cultures were gently agitated twice daily.

*S. trochoidea* was cultured until a density of 3,120 cells mL^−1^ was achieved on a modified f/2 medium without silicon. To avoid variable concentrations of Ca^2+^ from the initial seeding liquid, the cells were concentrated on a sterile Nitex screen, washed with sterile filtered (0.22μm pore size; Nucleopore filter) seawater, and resuspended in the modified experimental medium (Subba Rao 2011). A 50 mL sample of the culture was washed and inoculated into 500 mL of the f/2 culture medium in a 1,000 mL flask.

The cyst formation ratio (*S*) was calculated using the number of vegetative cells and cysts as follows:

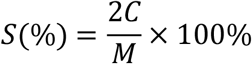

Where *M* is the maximum number of vegetative cells and *C* is the maximum number of cysts (Tomoyuki Shikata et al. 2008).

In this study, we examined the effects of Ca^2+^ on *S. trochoidea* growth by conducting five experiments in triplicate (Table 1). Of these five experiments, one group served as the control with no addition of CaCl_2_. In experimental groups two, three, four, and five, Ca^2+^ was added as CaCl_2_ at concentrations of 0.2, 0.4, 0.6, and 0.8 g·L^−1^, respectively. Given that there is often 0.4 g·L^−1^ of Ca^2+^ in seawater naturally, final Ca^2+^ concentrations in our experiments were 0.4, 0.6, 0.8, 1.0, and 1.2 g·L^−1^, respectively. Other required nutrients, trace metals, and vitamins were added to the f/2 medium and the cultures were homogenized by shaking before sampling.The duration of the experiment was 60 d. Samples were collected for cell counts daily at the start of the light cycle. Sample volumes of 3 mL were fixed with a drop of formalin, and resting cysts and motile cells were counted and photographed using an inverted microscope (Nikon, Ni-U, Tokyo, Japan).

**Tab.1.**
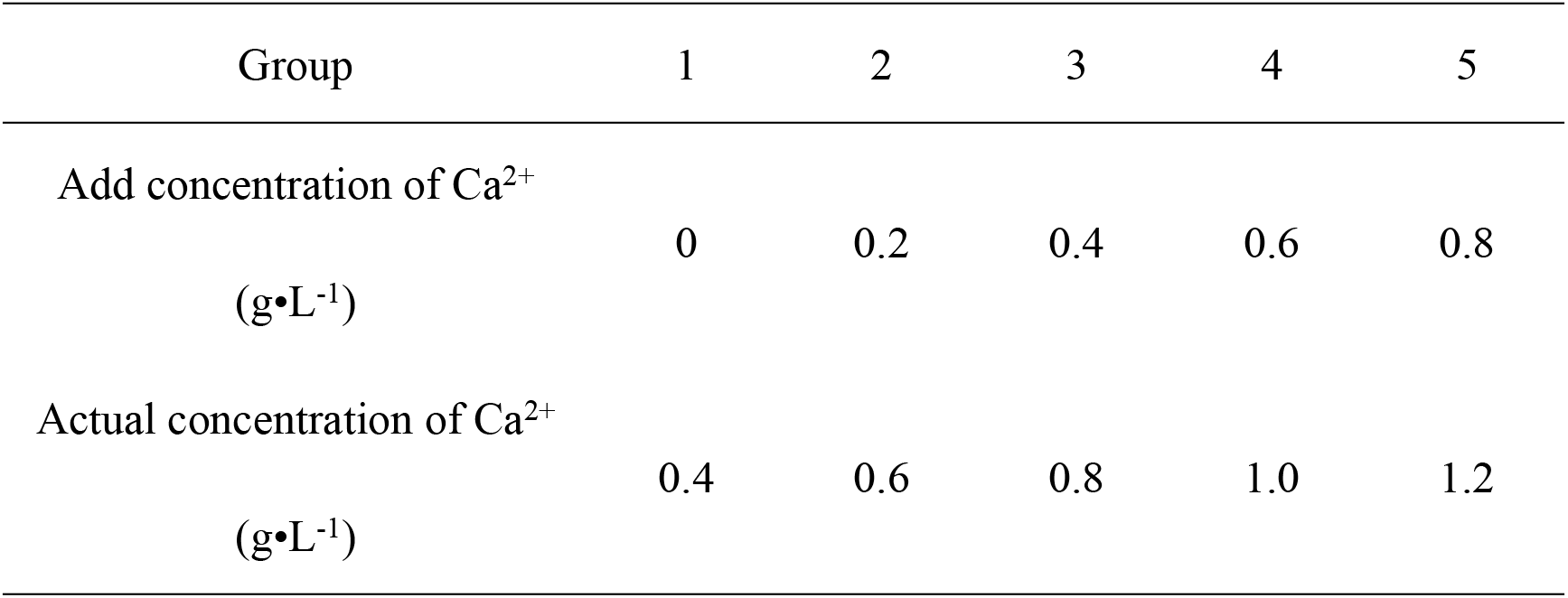
The different concentrations of Ca^2+^ in used in *S. trochoidea* culture experiments

## Results and Discussion

Resting cysts and motile cells were easily distinguishable in the present study. Motile cells were tapered and swam quickly, whereas resting cysts were egg or oval shaped, did not move, and two cyst shapes were predominantly observed: calcareous thorn cyst and smooth surface cysts.

### *S. trochoidea* motile cell growth under different Ca^2+^ treatments

We designed the laboratory experiments to study the response of *S. trochoidea* to different concentrations of Ca^2+^. Figure 1 shows changes in motile cell densities over time under different experimental conditions. Under different Ca^2+^ concentrations, the cell cultures did not enter a lag phase and cell numbers increased exponentially following starvation. *S. trochoidea* motile cell density increased as Ca^2+^ concentrations increased when the Ca^2+^ concentration was less than 0.2 g·mL^−1^. Cell density was then reduced as Ca^2+^ concentration exceeded 0.2 g·mL^−1^. As Ca^2+^ concentration increased, the stable and death phases were induced earlier in motile cells. The control group entered the stable phase on day 21 and death phase on day 35. Experimental groups exposed to Ca^2+^ concentrations of 0.2, 0.4, 0.6, and 0.8 g·L^−1^ entered stable phases on day 21, 18, 13, and 13, respectively, and entered death phases on day 35, 30, 30, and 27, respectively.

**Fig. 1.**
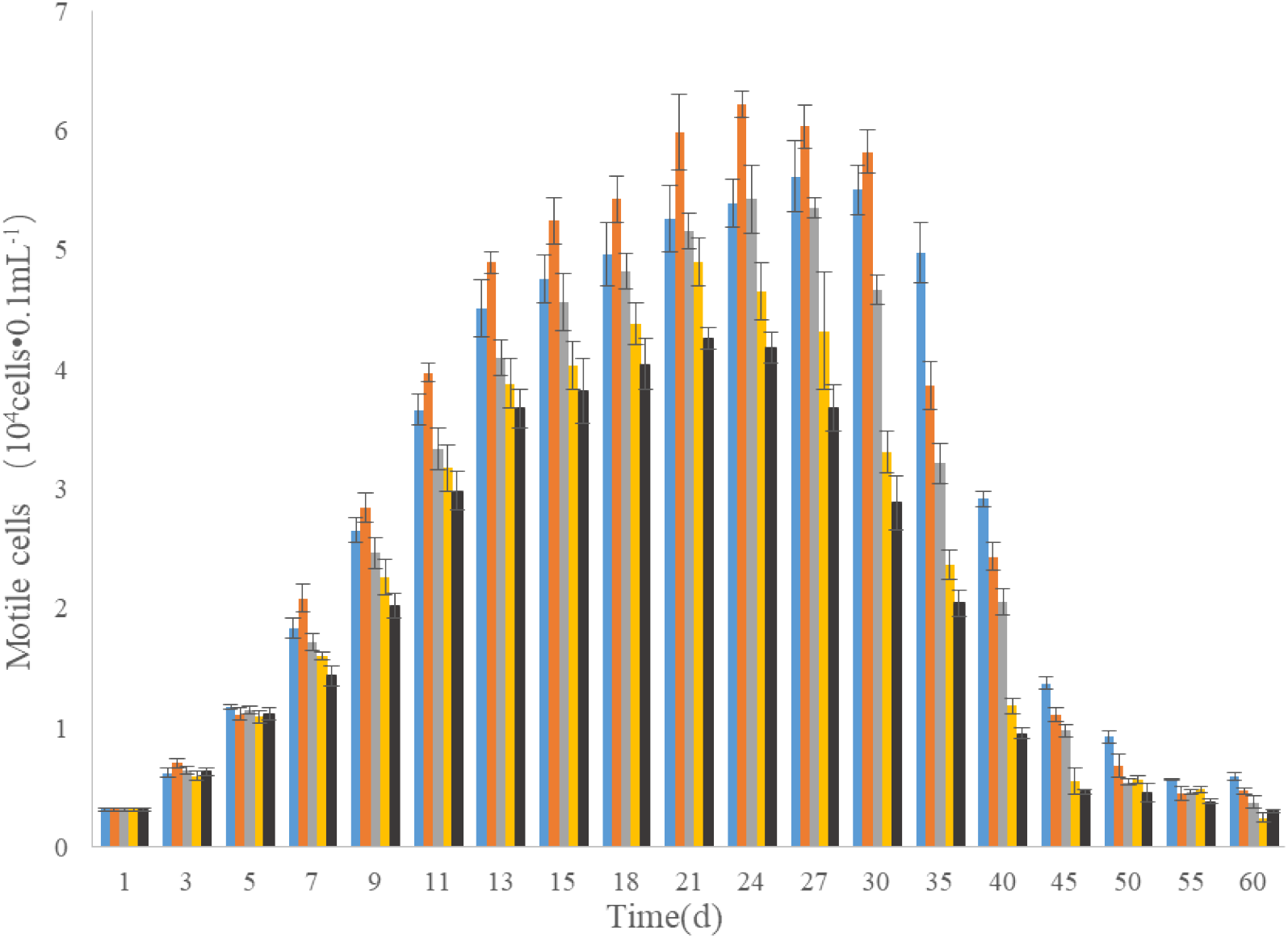
Changes in *S. trochoidea* motile cell densities over time under different Ca^2+^ treatment conditions 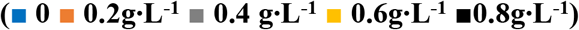

Some researchers studied the effects of Ca^2+^ on *Microcystis aeruginosa* growth and revealed that its growth was not influenced by increasing Ca^2+^ concentrations, although calcium was an important element for the growth of this species (Ding, 2017). However, there exist few studies which examined the effects of Ca^2+^ on the growth of different algal species and the results of these studies were variable. For example, Shi found that high concentrations of Ca^2+^ significantly inhibited the growth of *M. aeruginosa* (Shi et al. 2013). Alternatively, Li et al (2003) found that increased concentrations of Ca^2+^ in the culture medium stabilized the structure and function of *Anabaena* sp. PCC7120 cell membranes. Further, researchers revealed that *M. aeruginosa* growth was strongly inhibited by Ca^2+^ in that its growth decreased as the concentration of Ca^2+^ increased, but the effects of Ca^2+^ on *Scenedesmus obliquus* growth was less obvious(Zhao et al. 2014). Li et al (2017) further noted that increased Ca^2+^ concentrations promoted growth and improved biological calcification in *Microcystis flos-aquae*. Moreover, Huang (2012) found that both calcium and irradiance significantly influenced growth, colony formation, and colonial cell distribution in *Phaeocystis globose* in that growth and colony formation in this species were completely inhibited on calcium-free medium. Additionally, colony enlargement and abundance were hampered by low calcium concentrations. Lastly, compared with non-colony-forming cells, colony-forming cells favored high calcium conditions. Overall, these previous studies clearly reveal the variable influences of Ca^2+^ on different algal species.

### *S. trochoidea* cyst formation under different Ca^2+^ treatments

Figure 2 shows changes in *S. trochoidea* resting cyst densities over time under different experimental conditions. In the control group, cysts accumulated in large numbers after 15 d. In contrast with the experimental cultures, the control group formed cysts earlier and had a higher cyst density during early growth stages. The maximum cyst density(12.8×10^3^ cysts·mL^−1^) occurred in group three (0.4 g · L^−1^), and the minimum cyst density (6.2×10^3^ cysts·mL^−1^) occurred in the control group; this maximum cyst density was twice the minimum density. Cyst densities increased as Ca^2+^ concentrations increased until these concentrations reached 0.4 g·mL^−1^. Above this concentration, cyst densities began to decline with increasing Ca^2+^ concentrations.

**Fig. 2.**
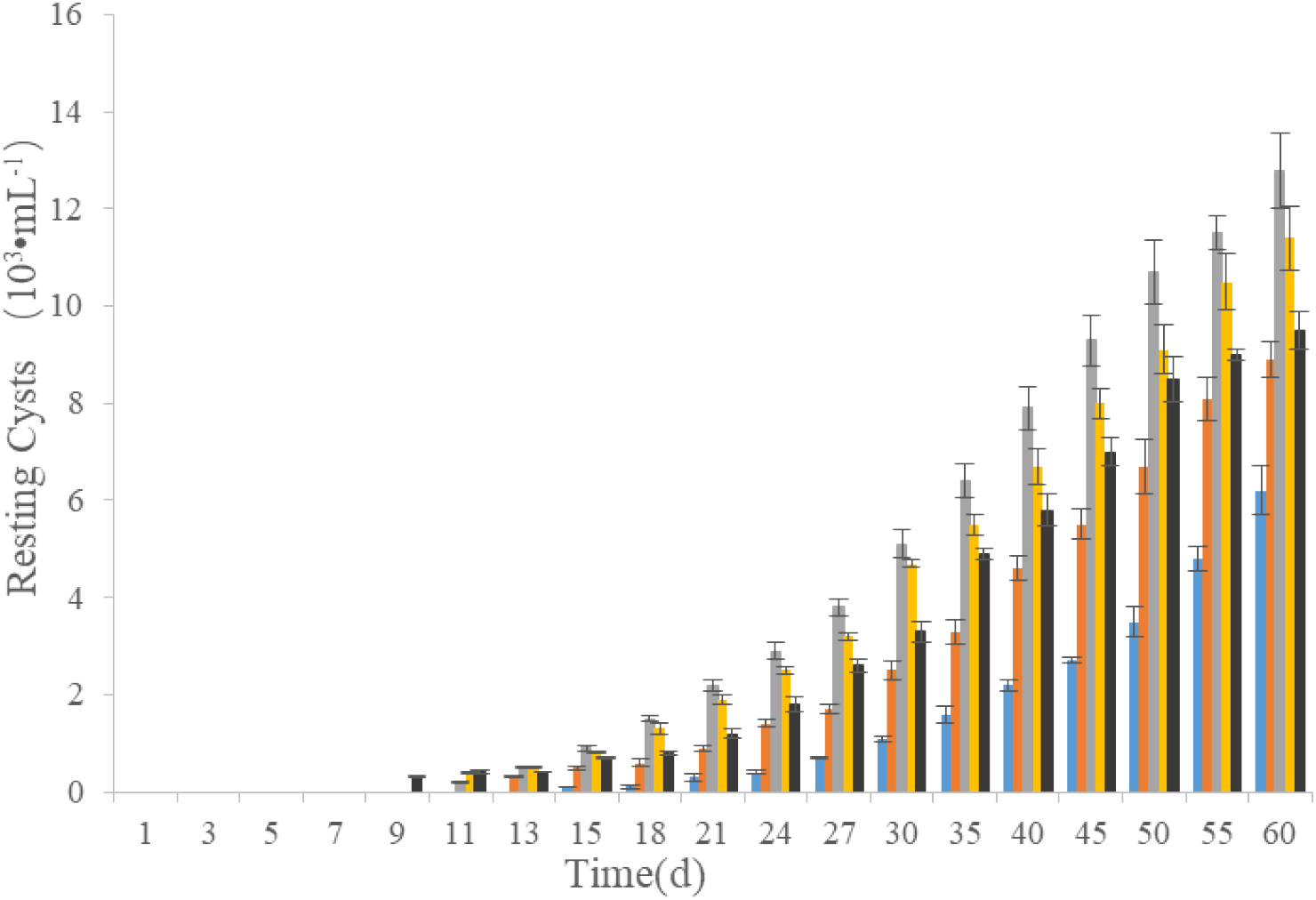
Changes in *S. trochoidea* cyst densities over time under different Ca^2+^ treatment conditions 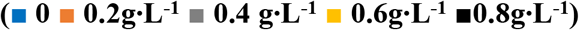

We further calculated the cyst formation ratio to examine the relationship between *S. trochoidea* cyst formation and varying Ca^2+^ concentrations (Figure 3). In this study, cyst formation ratios were 22.1%, 28.7%, 47.2%, 46.5 %, and 44.6% in *S. trochoidea* cultures exposed to Ca^2+^ concentrations of 0, 0.2, 0.4, 0.6, and 0.8 g·L^−1^, respectively. The highest cyst formation ratio was recorded in group three (0.4 g·L^−1^), and the lowest cyst formation ratio was recorded in the control group. These results indicate that cyst formation increased as Ca^2+^ concentration increased until these concentrations reached O.4g·L^−1^. Above this concentration, cyst formation was reduced as Ca^2+^ concentrations increased, but this reduction was not obvious.

**Fig. 3.**
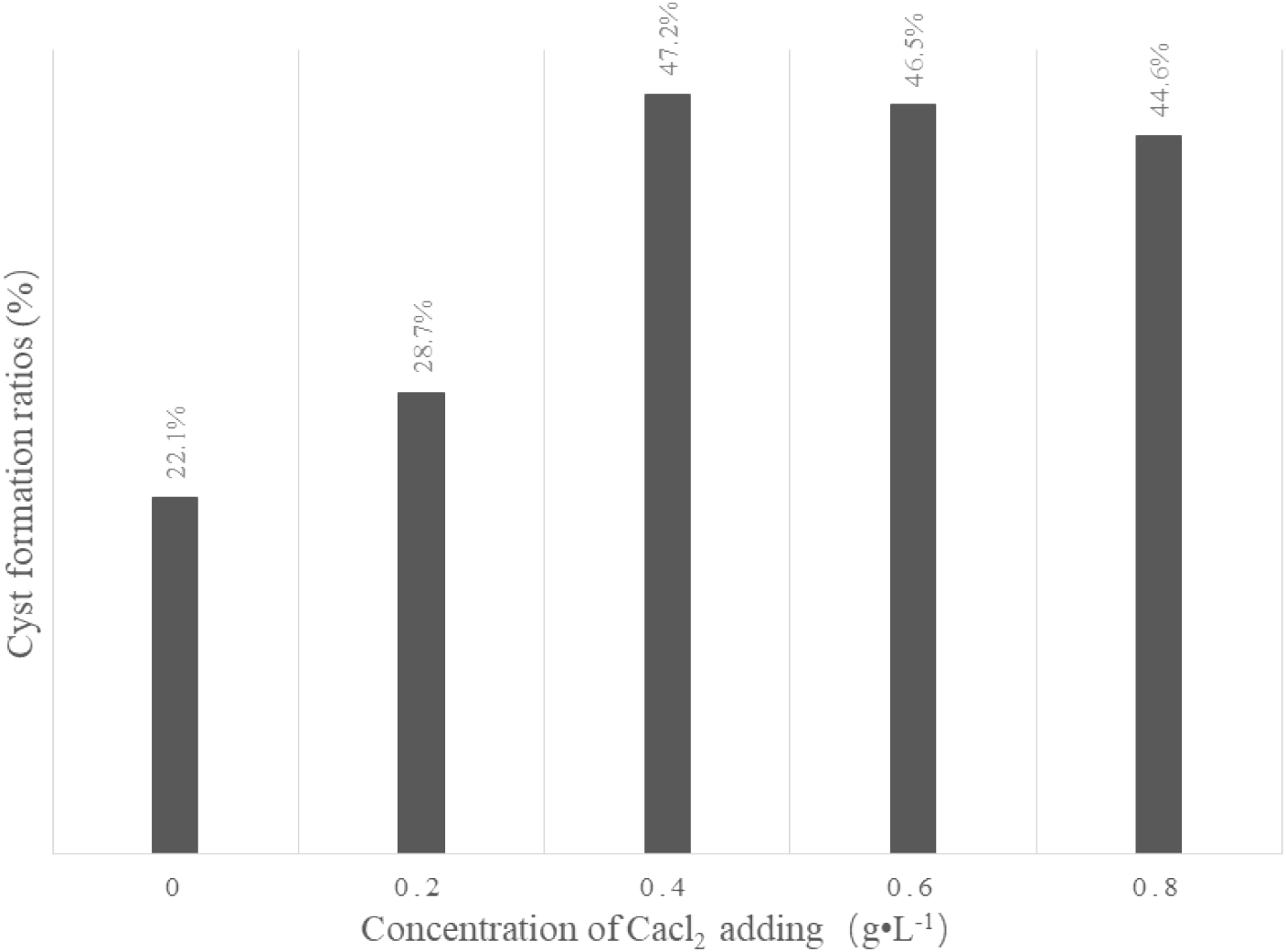
Cyst formation ratios in *S. trochoidea* cultures under different concentrations of CaCl_2_

To statistically analyze the effects of Ca^2+^ on encystment, we conducted a one-sample *T*-test (P < 0.01; SPSS 16.0) which revealed that Ca^2+^ addition significantly promoted cyst formation in *S. trochoidea* cell cultures (P = 0.004).

In the current study, high Ca^2+^ concentrations generally induced the formation of cysts in *S. trochoidea*. Moreover, these cysts appeared earlier at higher concentrations of Ca^2+^. For example, cysts were observed as soon as the ninth day in group 5 (treated with 0.8 g·L^−1^ of Ca^2+^). However, as Ca^2+^ concentrations increased, cyst formation rates initially increased and then decreased. This eventual decline in the rate of cyst formation may have occurred given that high concentrations of Ca^2+^ inhibited *S. trochoidea* growth in the current experiment. Lastly, it is important to note that on the last day of experimentation (60 days), the number and formation rate of cysts were less than the initial values.

### Rate of different cyst shapes in different concentrations of Ca^2+^

In our previous study, we identified two main *S. trochoidea* cyst shapes: calcareous thorn cysts and smooth surface cysts. In this study, we analyzed the formation rates of these two cysts under different Ca^2+^ concentrations. Figure 4 shows that the rate of calcareous thorn cyst formation increased as the concentration of Ca^2+^ increased, whereas the rate of smooth surface cyst formation showed an opposite trend. In the control group, calcareous thorn cysts reached high proportions (70.6%). However, in *S. trochoidea* cultures treated with Ca^2+^ concentrations of 0.2, 0.4, 0.6, and 0.8 g·L^−1^, the rates of calcareous thorn cyst formation were higher (75.0%, 80.8%, 81.5%, and 83.3%, respectively). Overall, our results indicated that under higher concentrations of Ca^2+^, more calcareous thorn cysts formed in *S. trochoidea* cultures.

**Fig. 4.**
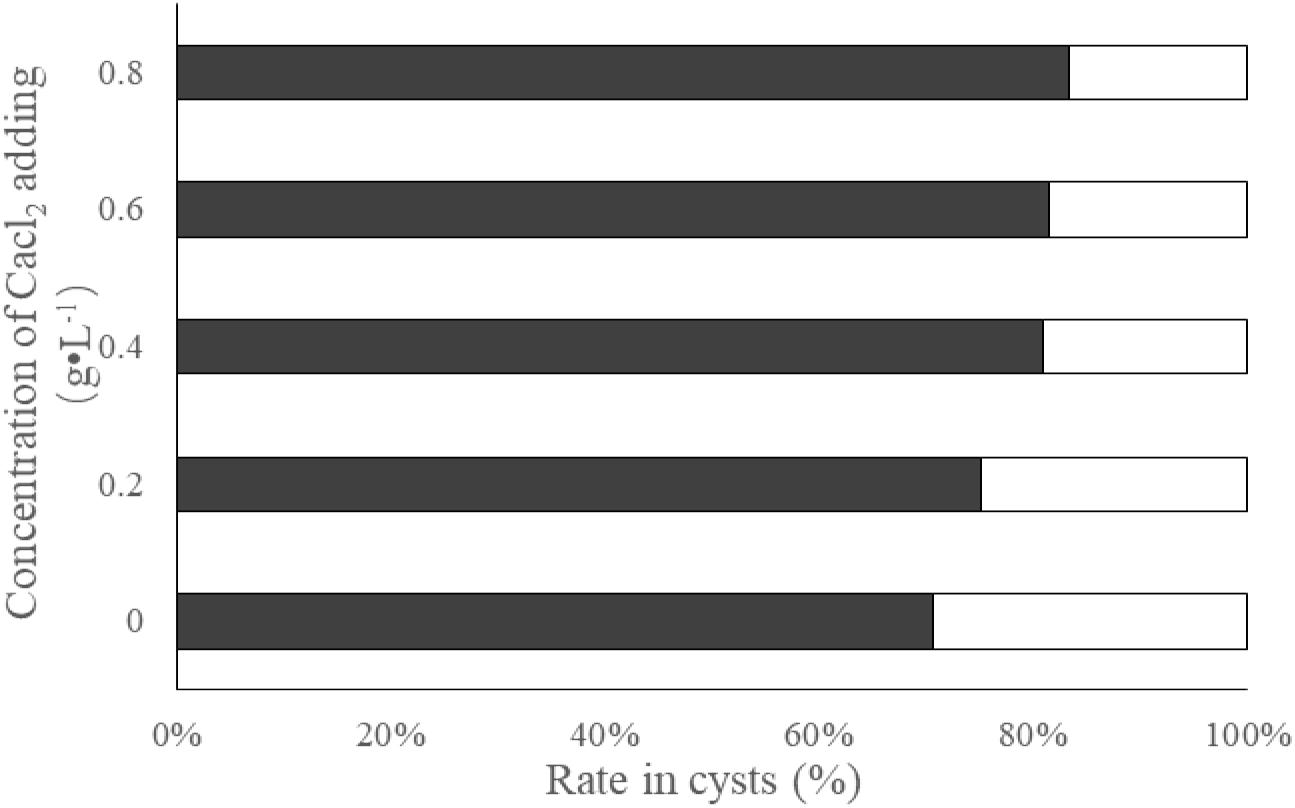
Rate of cyst formation in *S. trochoidea* cultures under different concentrations of Ca^2+^ on day 60 of experimentation (▪ Calcareous thorn cyst □ Smooth surface cyst)

To statistically analyze the effects of Ca^2+^ on the formation rate of these two cysts, we conducted a one-sample *T*-test (P < 0.05; SPSS 16.0) which revealed that Ca^2+^ addition significantly increased the rate of calcareous thorn cyst formation (P = 0.003).

Generally, calcareous thorn cysts were most abundant; however, the ratio of smooth surface cysts increased under some experimental conditions(Wang, 2014). In this study, the concentration of Ca^2+^ indeed influenced the ratio of these cysts in *S. trochoidea*. The ratio of calcareous thorn cysts increased with increasing concentrations of Ca^2+^, indicating that *S. trochoidea* absorbed Ca^2+^ from the water when cysts were formed, in accordance with previous research. For example, Wang Yan et al (2009) found that cyst formation was closely associated with Ca^2+^ in the water in that *S. trochoidea* absorbed Ca^2+^ to form the calcareous thorn following cyst formation. However, the increase in calcareous thorn cyst ratio gradually stabilized as Ca^2+^ concentrations increased. This result indicated that the shapes of cysts were controlled by the internal rhythms of *S. trochoidea*. In accordance with our previous studies, *S. trochoidea* indeed also formed smooth surface cysts although calcareous thorn cysts occurred at much higher ratios. This phenomenon requires further investigation, possibly at the genetic level.

## Conclusion

This study suggests that different concentrations of Ca^2+^ have different effects on growth and cyst formation in *S. trochoidea*. As Ca^2+^ concentrations increased, cell density initially increased and then decreased, and cell stable and death phases were induced earlier. The number of cysts and the cyst formation rate showed similar trends as vegetative cell density in response to increased Ca^2+^ concentrations in that they increased as Ca^2+^ concentration increased. After Ca^2+^ concentrations exceed 0.4 g·mL^−1^, the number of cysts and the cyst formation rate were reduced. We further revealed that *S. trochoidea* absorbed Ca^2+^ from the water when cysts were formed and higher Ca^2+^ concentrations promoted the formation of calcareous thorn cysts.

## Acknowledgments

This work was supported by Zhejiang Provincial Natural Science Foundation of China (Grant numbers LY15D060004) and the National Natural Science Foundation of China (Grant number 41776119).

## References

Tang YZ, Hu ZX, Deng YY. Characteristical life history (resting cyst) provides a mechanism for recurrence and geographic expansion of Harmful Algal Blooms of Dinoflagellates: a review. Studia marina sinica. 2016; 51:132–154.

Blackburn S. I, Parker N. Microalgal life cycles: encystment and excystment. In: Andersen RA (ed) Algal culturing techniques. Elsevier Academic Press, Amsterdam. 2005; 399–417.

Rosa Isabel Figueroa, Karin Rengefors, Isabel Bravo, Staffan Bensch. From homothally to heterothally: mating preferences and genetic variation within clones of the dinoflagellate *Gymnodinium catenatum*. Deep Sea Res Part II. 2010; 57:190–198.

Tang YZ, Gobler CJ. The toxic dinoflagellate *Cochlodinium Polykrikoides* (Dinophyceae) produces resting cysts. Harm Algae. 2012; 20:71–80.

Wang ZH, Qi YZ, Yang YF. Cyst formation: an important mechanism for the termination of *Scrippsiella trochoidea* (Dinophyceae) bloom. J Plankton Res. 2007; 29:209–218.

Ishikawa A, Taniguchi A. Contribution of benthic cysts to the population dynamics of *Scrippsiella* spp. (Dinophyceae) in Onagawa Bay, northeast Japan. Mar Ecol Prog Ser. 1996; 140:169–178.

Zhaohui Wang, Kazumi Matsuoka, Yuzao Qi, Jufang Chen, Songhui Lu. Dinoflagellate cyst records in recent sediments from Daya Bay, South China Sea. Phycol Res. 2004; 52:396–407.

Cho H J, Matsuoka K. Distribution of dinoflagellate cysts in surface sediments from the Yellow Sea and East China Sea. Mar Micropaleontol. 2001; 42:103–123.

Wang Zhifu, Yu Zhiming, Song Xiuxian, Cao Xihua, Han Xiaotian. Effects of modified clay on cysts of *Scrippsiella trochoidea*, Chinese Journal of Oceanology and Limnology. 2014; 32(6):1373–1382.

István Grigorszky, Kevi T. Kiss, Viktória Béres, István Bácsi, Márta M-Hamvas, Csaba Máthé, Gábor Vasas et al. The effect of temperature, nitrogen, and phosphorus on the encystment of *Peridinium cinctum*, Stein (Dinophyta). Hydrobiologia. 2006; 563: 527–535.

Subba Rao DV, Stewart JE. A preliminary study of the formation of a third category of cysts dinoflagellate, *Alexandrium fundyense* in response to elevated concentrations of ammonium chloride. Harmful Algae. 2011; 10: 512–52.

Tomoyuki Shikata, Sou Nagasoe, Tadashi Matsubara, Yasuhiro Yamasaki, Yohei Shimasaki. Encystment and Excystment of *Gyrodinium instriatum* Fredudenthal et Lee. Journal of Oceanography. 2008; 64:355–365.

Guillard R.R., Kilham, P., Jackson, T.A. Kinetics of Silicon-Limited Growth in Marine Diatom *Thalassiosira-Pseudonana* Hasle and *Heimdal Cyclotella-Nana* Hustedt. Journal of Phycology. 1973; 9(3), 233–237.

Shi J Q, Wu Z X, Song L R. Physiological and molecular responses to calcium supplementation in *Microcystis aeruginosa* (Cyanobacteria). New Zealand Journal of Marine and Freshwater Research. 2013; 47(1):1–11.

Ding Jiabo. Study on the impact of main nutrients as Ca > Mg on growth of *Microcystis aeruginosa*. Nanchang University, Marster, 2017.

Hao Li, Jinlai Miao, Fengxia Cui, Xiaoguang Liu, Guangyou Li. Effects of exogenous Ca^2+^ on the membrane permeability and photosynthetic characteristics of *Anabaena* sp. PCC7120 cells under simulated microgravity. Acta Hydrobiologica Sinica. 2003; 27(2): 136–139.

Zhao Lianfang, Ding Xiaoyan, Lu Lin, Li Ming. Effect of nutrient conditions an Ca^2+^ on the growth and competition of *Microcystis aeruginosa* and *Scenedesmus obliquus*. Environmental Science and Technology. 2014, 37(1): 13–17.

Li Xiaomin, Luo Kemeng, He Feng, Yang Zhenni, Fan Wenhong. Influence of calcium concentration on CO2 fixation with bio-calcification of *Microcystis flos-aquae*. Chinese Journal of Environmental Engineering. 2017; 11(12), 6327–6331.

Huang Tianwu. Morphological characteristics of colony for *Phaeocystis globosa* and the influences of light and Calcium Ion on formation and cell distribution of colony. Jinan University, Master, 2012.

Wang Yan, Xiong Defu, Gu Haifeng, Li Shaoshan. Morphological characters of *Scrippsiella trochoidea* cysts. Chinese Bulletin of Botany. 2009; 44(6): 701–709.

